# Observing Aggression Increases Aggression In Semi-Free Ranging Barbary Macaques

**DOI:** 10.1101/2023.01.12.523737

**Authors:** Rachel A. Blood, Stuart Semple

## Abstract

In many social living species, seeing conspecifics interacting can alter the behaviour of bystanders, leading to social contagion – the spread of behaviour or emotion among group members. Among primates, studies of a small number of species have explored bystanders’ responses to observing aggressive interactions, finding evidence that individuals that see such interactions are more likely to subsequently engage in aggressive behaviours themselves. To increase the taxonomic breadth of this body of research, working with semi-free ranging adult female Barbary macaques (*Macaca sylvanus*) at the Trentham Monkey Forest, Stoke-on-Trent, UK, we assessed bystanders’ responses to observing naturally occurring aggressive interactions. Data were collected under two conditions: (i) after observing an aggressive interaction between conspecifics and (ii) during a matched-control period, before which individuals did not observe aggression. Bystanders were significantly quicker to initiate an aggressive interaction themselves after observing an aggressive interaction than if they had not, providing evidence of behavioural contagion. There was no effect of observing aggression on the rates of self-directed behaviour (an indicator of anxiety), suggesting the negative emotional states associated with involvement in aggressive interactions did not spread to bystanders. The results of this study provide new insight into the nature and importance of visual contagion of behaviour among primates.

## Introduction

The spread of emotion or behaviour from one individual to others is known as social contagion[1–3]. This phenomenon can promote group cohesion [4], behavioural synchronization [5], social bonding [6], and social learning opportunities [4] among individuals within group living-species. Understanding the nature and extent of social contagion is a key goal for many animal behaviour researchers, because of the possible impacts of this phenomenon on individuals and their social networks [3]. Most studies in this area tend to focus on visual contagion – the spread of a behaviour via visual observation – which has been documented among a range of non-human animal species.

Such studies often look specifically at self-directed behaviours, i.e., self-scratching [7] or yawning [8], and the spread of such behaviours has been documented among a diverse range of social species including rats, *Rattus norvegicus* [15], wolves, *Canis lupus lupus* [16] and a range of primates - orangutans, *Pongo-pygmaeus* [7], chimpanzees, *Pan troglodytes* [17–18], geladas, *Theropithecus gelada* [19], and rhesus macaques, *Macaca mulatta* [14]. Social contagion can occur not only when bystanders observe the behaviour of another individual animal, but also when attending to dyadic social interactions [1–2, 20]. Studies of bystanders’ response to both positive (i.e., affiliative) and negative (i.e., agonistic) social behaviours have provided evidence of this form of contagion.

Evidence for contagion following perception of positive interactions has been found in a range of primate species. Captive common marmosets, *Callithrix jacchus*, that were shown videos of conspecifics grooming exhibited increased rates of grooming behaviours themselves [2]. Among semi-free ranging Barbary macaques, *Macaca sylvanus* [22] and rhesus macaques, *Macaca mulatta* [23], bystanders that observed others grooming were themselves subsequently quicker to initiate grooming interactions. In the Barbary macaque study, individuals also showed reduced rates of self-directed behaviours after observing grooming, suggesting a reduction in anxiety levels; this indicates that witnessing affiliative interactions can, in addition to behavioural contagion, lead to a spread of the associated emotional effects [22].

Visual contagion linked to negative social interactions has also been documented in a range of primates. In wild vervet monkeys, *Chlorocebus pygerythrus* [24] and captive Japanese macaques, *Macaca fuscata* [25], the kin of victims of an aggression frequently show aggression shortly afterwards towards the kin of the original aggressor. In captive mandrills, *Mandrillus sphinx*, bystanders to aggressive interactions themselves subsequently show increased rates of aggression [26], although here this was not directed particularly at kin of the aggressor observed. As aggression is a ubiquitous and frequently occurring social behaviour among gregarious species [27–29], adding to the limited body of research on its possible spread through visual observation will provide greater insight into the nature and impact of social contagion.

In this study, we test the hypothesis that there is visual contagion of aggression in Barbary macaques, predicting that observing an aggressive interaction will reduce the time for bystanders to initiate an aggressive interaction themselves (*Prediction 1*) Additionally, we assess whether observing aggressive behaviour is associated with negative changes in an individual’s affective state, predicting that observing an aggressive interaction will increase rates of self-directed behaviour among bystanders (*Prediction 2*).

## Methods

### Study Site and Subjects

We conducted this study with the semi-free ranging Barbary macaques at Trentham Monkey Forest (Stoke-on-Trent, UK). This site is home to approximately 140 individuals across three social groups, in a fenced-in 24 ha area of grassland and cedar and oak forest. The park is open daily to visitors who can only walk on designated paths and may not feed or touch the animals. The macaques are provisioned by staff several times a day with a mixture of fruits, vegetables, seeds, and grains. Each macaque has a unique code tattooed on the inside of their thigh for individual identification. The subjects of this study were 26 adult females from the same social group, with ages ranging from 6 to 28 years of age. Their social group was comprised of 53 individuals at the start of the study: 18 adult males (older than 8 years of age), 26 adult females (older than 6 years of age), 5 subadult individuals (males less than 8 years of age and females less than 6 years of age), and 4 juveniles (less than 3 years of age). During the study period, six of the study females gave birth. Data were collected on those pregnant females up to the birth of their offspring, but not afterwards.

### Data Collection

We conducted behavioural observations between 0900 and 1700 from 28 March to 28 June 2022. All behavioural observations were recorded using an iPod Touch equipped with the ANIMAL BEHAVIOUR PRO© v. 1.2 application [30]. We collected the data using an adapted version of the well-established post-conflict/matched-control (PC-MC) method [31]. Used originally to study post-conflict behaviour, this procedure allows for observational data to be collected on individuals after they had been involved in a conflict (PC) and then compared to data collected on the same individual in the absence of any conflict (MC). In this study, following similar modifications to this method by Berthier and Semple [22] and Ostner et al. [23], we collected data on bystander individuals after they observed an aggressive interaction as well as in a matched control period.

### After Observing an Aggressive Interaction

We took ‘After observing an aggressive interaction’ (AOAI) samples opportunistically. The samples began when an aggressive interaction (a lunge, hit, grab, bite, chase, or threat facial expression) started between two individuals and there was a study adult female as a bystander within 7 m, that had their eyes open and that was oriented in the general direction of the interaction at least once. For each AOAI, we recorded the age class/sex of the aggressor and victim of the aggressive interaction as well as the nature of their interaction. The number, age class and sex of animals within 10m of the focal individual and the proximity of the nearest animal during the observation of an aggressive interaction were also noted, to ensure the best consistency with the matched-control focal. We then followed the female bystander in a continuous focal watch, which ended when they themselves initiated an aggressive interaction. During this time, we recorded the time of occurrence and nature of the first aggressive interaction initiated by the focal individual, as well as all occurrences of self-directed behaviours (self-grooming, self-scratching, yawning). If the focal individual did not initiate aggression within two hours of the start of the AOAI sample, the observation was stopped.

### Matched-Control Period

On the next available day following the AOAI, we carried out a corresponding matched-control observation (MC), starting at about the same time of day as the corresponding AOAI (+/− 60 minutes), collecting the same type of data (all occurrences of self-directed behaviours, and the time of occurrence and nature of the first aggressive interaction initiated by the focal individual) during the same amount of time as the corresponding AOAI or until the focal individual initiated an aggressive interaction, whichever came first. As best as possible, we matched the social environment of the MC and AOAI - i.e., number of animals and their age class and sex within 10m of the focal, and the proximity of nearest animal. We monitored the focal individual for a 10-minute period before the beginning of the MC period to ensure that they were not a bystander to an aggressive interaction in this time.

### Data Analyses

To test Prediction 1, following de Waal and Yoshihara [31], AOAI – MC pairs were classified as ‘attracted’ (when the focal individual initiated an aggressive interaction in the AOAI period but not in the corresponding MC period), ‘dispersed’ (when the focal individual initiated an aggressive interaction before the end of the MC period, including the AOAI – MC pairs where aggression was not initiated in the AOAI period but did occur in the MC period), or ‘neutral’ (aggression was not initiated in either the AOAI or MC periods). To test Prediction 2, for rates of SDB per minute, AOAI – MC pairs were classified as ‘higher in AOAI’ (when rates of SDB were greater in the AOAI period than in the MC period), ‘lower in AOAI’ (when rates of SDB were lower in the AOAI period than in the MC period), and ‘neutral’ (when rates of SDB were equal across a AOAI – MC pair) for each AOAI – MC pair collected.

Subsequent analyses were conducted to check whether AOAI and MC pairs differed in relation to number and closest proximity of conspecifics. For the number of neighbours within 10 m of the focal individual at the start of an observation, AOAI – MC pairs were classified as ‘greater in AOAI’ (when the number of neighbours was greater in the AOAI period than in the MC period), ‘fewer in AOAI’ (when the number of neighbours was lower in the AOAI period than in the MC period), and ‘neutral’ (when the number of neighbours was equal across a AOAI – MC pair) for each AOAI – MC pair collected. for distance to the nearest neighbour at the start of the focal, AOAI – MC pairs were classified as ‘further in AOAI’ (when the nearest neighbour was further away from the focal individual in the AOAI period than in the MC period), ‘closer in AOAI’ (when the nearest neighbour was closer to the focal individual in the AOAI period than in the MC period), and ‘neutral’ (when the nearest neighbour was the same distance to the focal individual across a AOAI – MC pair) for each AOAI – MC pair collected.

To avoid pseudo replication, and following previously adopted methods in the field [22], for each study female (n=26) the proportion of ‘attracted’ and ‘dispersed’ AOAI – MC pairs and the proportion of ‘higher in AOAI’ and ‘lower in AOAI’ SDB rates across their AOAI – MC pairs were calculated. We used Wilcoxon matched-pairs tests to determine whether, over the 26 study females, the proportion of ‘attracted’ AOAI-MC pairs was greater than the proportion of ‘dispersed’ AOAI-MC pairs (*Prediction 1*), and whether the proportion of ‘higher in AOAI’ AOAI – MC pairs was greater than the proportion of ‘lower in AOAI’ AOAI – MC pairs (*Prediction 2*). To test for comparability between AOAI and MC pairs in terms of focal females’ social environments, the same approach was used to determine whether there was a significant difference between the proportion of ‘greater in AOAI’ and ‘fewer in AOAI’ AOAI – MC pairs, and between the proportion of ‘further in AOAI’ and ‘closer in AOAI’ AOAI – MC pairs. We conducted all tests in R version 4.2.1 [32].

## Results

A total of 126 AOAI – MC pairs were collected over the 26 adult females, which represents a total of 88 h 8 min of observation. The number of pairs per individual female ranged from 3 – 9. For 23 of the 126 AOAIs, the focal individual did not initiate an aggressive interaction; for 16 of these 23 AOAIs, aggression was not seen in the corresponding MC focal.

In support of *Prediction 1*, the average proportion of ‘attracted’ AOAI – MC pairs was higher than the average proportion of ‘dispersed’ pairs (attracted: median = 0.63, range = 0.00 – 1.00; dispersed: median = 0.20, range = 0.00 – 0.60; Wilcoxon matched-pairs test: Z = 3.907, n = 26, p <0.001; Figure 1). *Prediction 2* was not supported, as there was not a significant difference between the average proportion of ‘higher in AOAI’ AOAI – MC pairs and ‘lower in AOAI’ pairs (Z = 1.138, n = 26, p = 0.255; Figure 2). There was a large amount of ‘neutral’ AOAI – MC pairs for SDB (27 of the 126 pairs), mainly due to there not being SDB present in either the AOAI or the MC period.

**Figure 1.**
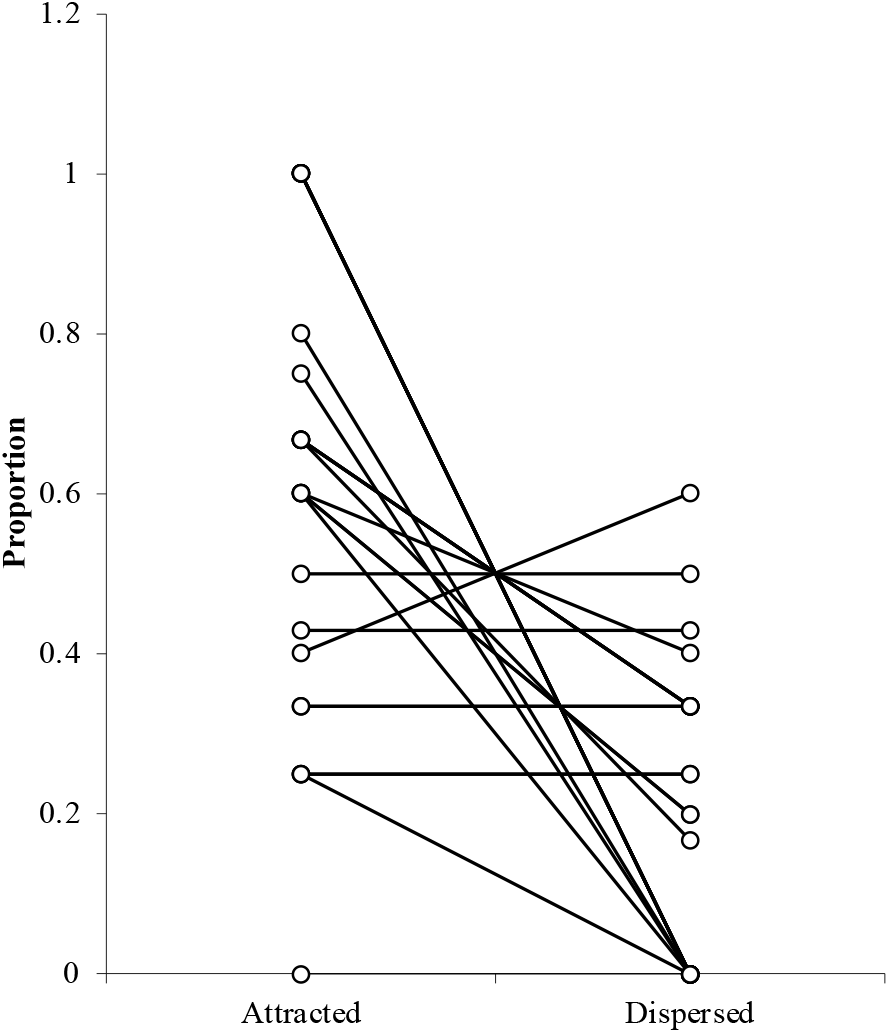
Proportion of attracted and dispersed ‘after observing an aggressive interaction (AOAI)’ and ‘matched control (MC)’ pairs. The lines connect data points for each individual female (n = 26). AOAI – MC pairs could also be neutral, so some values for individual females do not total 1. Certain lines overlap entirely, so the figure appears to have less than 26 lines.

**Figure 2.**
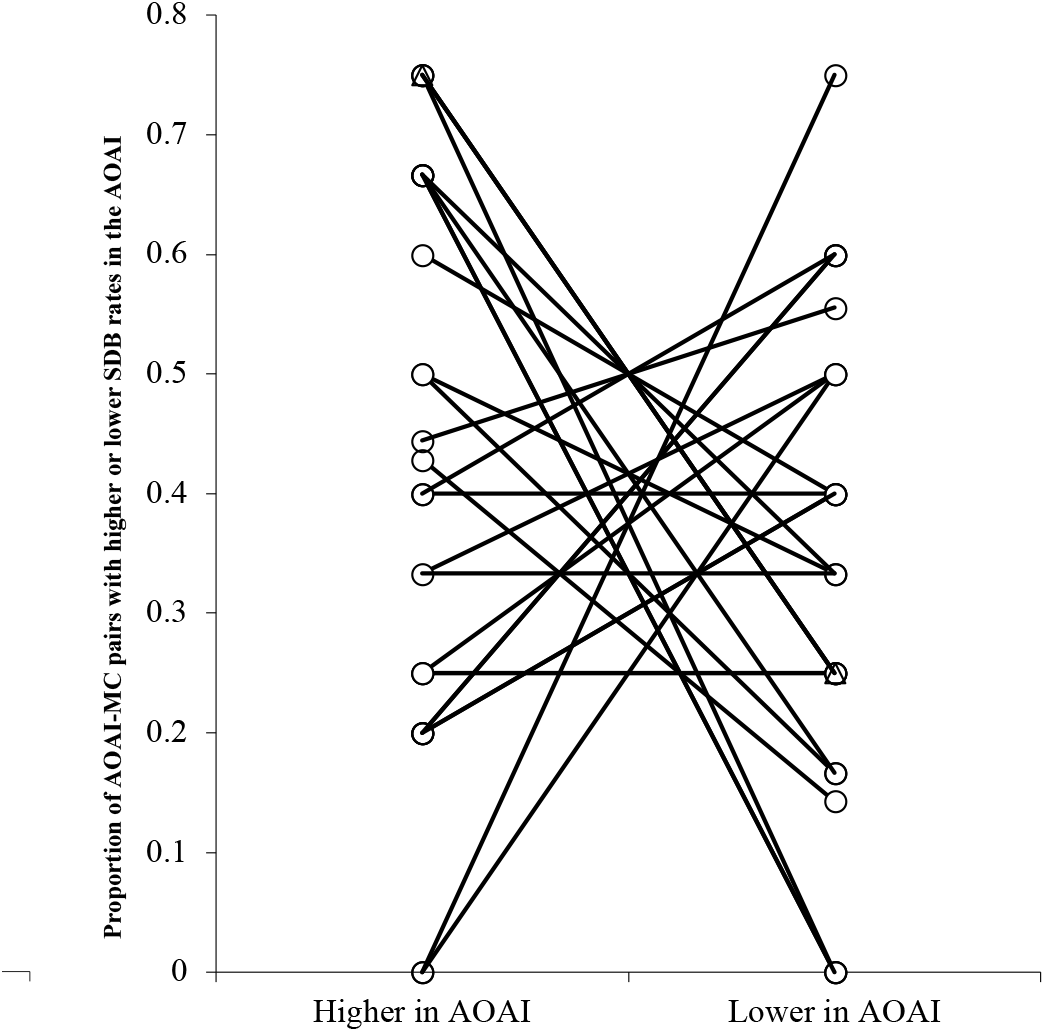
Proportion of aggressive interaction (AOAI) and matched control (MC) pairs in which rates of self-directed behaviours are ‘higher in AOAI’ or ‘lower in AOAI’. The lines connect data points for each individual female (n = 26). AOAI – MC pairs could also be neutral, so some values for individual females do not total 1. Certain lines overlap entirely, so the figure appears to have less than 26 lines.

There was not a significant difference in the number of neighbours within 10m of the individual at the start the focal (Z = 1.073, n = 26, p = 0.281). However, there was a greater proportion of AOAI – MC pairs where the nearest neighbour was further away from the focal individual in the AOAI than in the MC (Z = 2.391, n = 26, p = 0.017).

## Discussion

In this study of adult female Barbary macaques, we tested whether observing aggression leads to negative behavioural contagion - a spread of aggression - among group members. As predicted under this phenomenon, seeing conspecifics engage in aggressive interactions was associated with a decreased latency to subsequent aggression among bystanders. This finding adds a new primate species to the small number in which this phenomenon has been documented. Contrary to prediction, observing aggressive interactions was not associated with changes in bystanders’ SDB rates, indicating they did not experience an increase in levels of anxiety in such circumstances. Overall, this work further highlights the importance of understanding how social interactions can affect more than just the animals directly involved.

The evidence provided here for visual contagion of aggression suggests that agonistic interactions in this species have effects that reach well beyond those directly involved - aggression spreading from the original interaction partners to bystanders, from those bystanders to their new interaction partners, and potentially then also to the new bystanders to those interactions and beyond. The nature and extent of visual contagion effects may play an important role in shaping patterns of aggression within and between populations of primate species, and their variation between species may underpin inter-specific variation in social style [22]. In relation to the latter point, it is notable that Barbary macaques are classified as a ‘tolerant’ species, while similar contagion of aggression has been documented in the ‘despotic’ Japanese macaque [25]. An important goal for future work will be to develop methods that allow for more nuanced quantification of contagion effects - thus facilitating comparisons between species - rather than merely documenting their presence or absence.

The question of why visual contagion of aggression occurs, and in particular whether or not this is adaptive, is an important one. There is evidence that female Barbary macaques may use aggression as a coping strategy [39], and thus elevated rates of agonism among bystanders may reflect an adaptive response that mediates the emotional impacts of witnessing aggression. Bystanders may also benefit from their own aggression by pre-emptively decreasing the likelihood of being the victims of aggression themselves [27]. Alternatively, as argued by Schino and Sciarretta [26], bystanders’ increased tendency to aggression may be nonadaptive, resulting from broader social facilitation processes; under this explanation, contagion of aggression is a by-product of an adaptive general mechanism that evolved to promote behavioural synchronization between individuals, and thus cohesion at the group level.

Contrary to our prediction, female Barbary macaques did not show higher rates of SDBs after observing aggression than in matched control periods. This is in contrast to results from studies of a range of primate species [e.g., 26, 36], including wild male and female Barbary macaques [40], where observing aggressive interactions elevates SDB rates. An exception to this general pattern – presenting results in line with ours of no elevation in SDB among bystanders – is provided by a study of captive geladas, *Theropithecus gelada* [41]. The authors of that study linked this apparent lack of anxiety among bystanders to the species’ relatively tolerant social style, and as Barbary macaques also show such a social style this may be an explanatory factor. Alternatively, or in addition, females’ use of aggression as a coping behaviour [39] may underpin our results, with bystanders effectively mediating their anxiety through their own agonism such that SDB levels do not rise significantly.

One methodological finding also merits discussion – despite our best efforts to match focal females’ immediate social environment between AOAI and MC pairs, it was found that overall the nearest neighbour was further away from the focal individual in the AOAI than in the MC. We do not see an obvious way in which this difference might bias our findings, particularly as no such difference was seen for the number of conspecifics within 10m, but we cannot of course exclude that possibility. At the least, this analysis highlights the value of actually testing for differences between AOIA and MC pairs (or more traditional PC-MC pairs), rather than assuming – as has typically been done – that careful study execution eliminated the potential for such differences.

The findings of our study add to a growing body of work that highlights how aggression can spread though visual contagion. To understand this phenomenon more fully, it would be valuable to explore how factors such as sex, rank, age, and relatedness to the animals being observed may diminish or intensify the occurrence of the contagion. Identifying the full extent of the chain of contagion is another key goal. Perhaps most importantly, testing adaptive and non-adaptive hypotheses about visual contagion of aggression is needed to help understand its possible proximate and ultimate causes. Addressing these challenges will be critical if we are to have most complete understanding of the nature and evolution of this important social phenomenon.

## Ethics

We received ethics approval from the University of Roehampton, and Trentham Monkey Forest gave permission to conduct this study. We attempted to always keep a minimum distance of 3 m from all monkeys, and strictly avoided direct eye and physical contact with them.

## Authors’ Contributions

R.B. and S.S. designed the study and wrote the paper; R.B. collected and analysed the data.

## Competing Interests

We report no competing interests.

## Funding

This project was partially funded by the University of Roehampton.

## Acknowledgements

We would like to thank Trentham Monkey Forest for permission to conduct this study, and the Monkey Forest staff - especially Anna Smith and Diane Floyd - for their wonderfully generous assistance throughout data collection. We also thank Dr Harry Marshall for his helpful comments in the developmental stage of this study, and for his statistical advice.

## Notes

### Competing Interest Statement

The authors have declared no competing interest.

## References

1. Levy DA, Nail PR. Contagion: a theoretical and empirical review and reconceptualization. Genetic, social, and general psychology monographs. 1993 May.

2. Watson CF, Caldwell CA. Neighbor effects in marmosets: social contagion of agonism and affiliation in captive Callithrix jacchus. American Journal of Primatology: Official Journal of the American Society of Primatologists. 2010 Jun;72(6):549–58.

3. Christakis NA, Fowler JH. Social contagion theory: examining dynamic social networks and human behavior. Statistics in medicine. 2013 Feb 20;32(4):556–77.

4. Coussi-Korbel S, Fragaszy DM. On the relation between social dynamics and social learning. Animal behaviour. 1995 Jan 1;50(6):1441–53.

5. Gordon I, Gilboa A, Cohen S, Milstein N, Haimovich N, Pinhasi S, Siegman S. Physiological and behavioral synchrony predict group cohesion and performance. Scientific reports. 2020 May 21;10(1):1–2.

6. Launay J, Tarr B, Dunbar RI. Synchrony as an adaptive mechanism for large-scale human social bonding. Ethology. 2016 Oct;122(10):779–89.

7. Laméris DW, van Berlo E, Sterck EH, Bionda T, Kret ME. Low relationship quality predicts scratch contagion during tense situations in orangutans (Pongo pygmaeus). American journal of primatology. 2020 Jul;82(7):e23138.

8. Guggisberg AG, Mathis J, Schnider A, Hess CW. Why do we yawn?. Neuroscience & Biobehavioral Reviews. 2010 Jul 1;34(8):1267–76.

9. Maestripieri D. Maternal anxiety in rhesus macaques (Macaca mulatta) II. Emotional bases of individual differences in mothering style. Ethology. 1993 Jan 12;95(1):32–42.

10. Schino G, Perretta G, Taglioni AM, Monaco V, Troisi A. Primate displacement activities as an ethopharmacological model of anxiety. Anxiety. 1996;2(4):186–91.

11. Aureli F. Post-conflict anxiety in nonhuman primates: The mediating role of emotion in conflict resolution. Aggressive Behavior: Official Journal of the International Society for Research on Aggression. 1997;23(5):315–28.

12. Schino G. Grooming and agonistic support: a meta-analysis of primate reciprocal altruism. Behavioral Ecology. 2007 Jan 1;18(1):115–20.

13. Cheney DL, Seyfarth RM. Stress and coping mechanisms in female primates. Advances in the Study of Behavior. 2009 Jan 1;39:1–44.

14. Feneran AN, O’donnell R, Press A, Yosipovitch G, Cline M, Dugan G, Papoiu AD, NATTkEMPER LA, Chan YH, Shively CA. Monkey see, monkey do: contagious itch in nonhuman primates. Acta dermato-venereologica. 2013 Jan 1;93(1):27–9.

15. Moyaho A, Rivas-Zamudio X, Ugarte A, Eguibar JR, Valencia J. Smell facilitates auditory contagious yawning in stranger rats. Animal cognition. 2015 Jan;18(1):279–90.

16. Romero T, Ito M, Saito A, Hasegawa T. Social modulation of contagious yawning in wolves. PloS One. 2014 Aug 27;9(8):e105963.

17. Anderson JR, Myowa–Yamakoshi M, Matsuzawa T. Contagious yawning in chimpanzees. Proceedings of the Royal Society of London. Series B: Biological Sciences. 2004 Dec 7;271(suppl_6):S468–70.

18. Campbell MW, Carter JD, Proctor D, Eisenberg ML, de Waal FB. Computer animations stimulate contagious yawning in chimpanzees. Proceedings of the Royal Society B: Biological Sciences. 2009 Dec 7;276(1676):4255–9.

19. Palagi E, Leone A, Mancini G, Ferrari PF. Contagious yawning in gelada baboons as a possible expression of empathy. Proceedings of the National Academy of Sciences. 2009 Nov 17;106(46):19262–7.

20. Schülke O, Dumdey N, Ostner J. Selective attention for affiliative and agonistic interactions of dominants and close affiliates in macaques. Scientific reports. 2020 Apr 6;10(1):1–8.

21. Videan EN, Fritz J, Schwandt ML, Smith HF, Howell S. Controllability in environmental enrichment for captive chimpanzees (Pan troglodytes). Journal of Applied Animal Welfare Science. 2005 Apr 1;8(2):117–30.

22. Berthier JM, Semple S. Observing grooming promotes affiliation in Barbary macaques. Proceedings of the Royal Society B. 2018 Dec 19;285(1893):20181964.

23. Ostner J, Wilken J, Schülke O. Social contagion of affiliation in female macaques. Royal Society open science. 2021 Jan 13;8(1):201538.

24. Cheney DL, Seyfarth RM. The recognition of social alliances by vervet monkeys. Animal behaviour. 1986 Dec 1;34(6):1722–31.

25. Aureli F, Cozzolino R, Cordischi C, Scucchi S. Kin-oriented redirection among Japanese macaques: an expression of a revenge system?. Animal Behaviour. 1992 Aug 1;44:283–91.

26. Schino G, Sciarretta M. Effects of aggression on interactions between uninvolved bystanders in mandrills. Animal Behaviour. 2015 Feb 1;100:16–21.

27. Bernstein IS, Gordon TP. The function of aggression in primate societies: Uncontrolled aggression may threaten human survival, but aggression may be vital to the establishment and regulation of primate societies and sociality. American Scientist. 1974 May 1;62(3):304–11.

28. Lorenz K, Latzke M, Salzen E. On aggression. Routledge; 2021 Sep 23.

29. Hinde RA. The bases of aggression in animals. Journal of psychosomatic research. 1969 Sep 1;13(3):213–9.

30. Newton-Fisher NE. 2012 Animal Behaviour Pro: v. 1.4. 4.

31. De Waal FB, Yoshihara D. Reconciliation and redirected affection in rhesus monkeys. Behaviour. 1983 Jan 1;85(3-4):224–41.

32. R Development Core Team. 2022 The R project for statistical computing.

33. Tinbergen N. On War and Peace in Animals and Man: An ethologist’s approach to the biology of aggression. Science. 1968 Jun 28;160(3835):1411–8.

34. Honess PE, Marin CM. Behavioural and physiological aspects of stress and aggression in nonhuman primates. Neuroscience & Biobehavioral Reviews. 2006 Jan 1;30(3):390–412.

35. Barsade SG. The ripple effect: Emotional contagion and its influence on group behavior. Administrative science quarterly. 2002 Dec;47(4):644–75.

36. Judge PG, Mullen SH. Quadratic postconflict affiliation among bystanders in a hamadryas baboon group. Animal Behaviour. 2005 Jun 1;69(6):1345–55.

37. Majolo B, Ventura R, Koyama NF. Anxiety level predicts post-conflict behaviour in wild Japanese macaques (Macaca fuscata yakui). Ethology. 2009 Oct;115(10):986–95.

38. Paschek N, Müller N, Heistermann M, Ostner J, Schülke O. Subtypes of aggression and their relation to anxiety in Barbary macaques. Aggressive behavior. 2019 Mar;45(2):120–8.

39. Gustison ML, MacLarnon A, Wiper S, Semple S. An experimental study of behavioural coping strategies in free-ranging female Barbary macaques (Macaca sylvanus). Stress. 2012 Nov 1;15(6):608–17.

40. McFarland R, Majolo B. Reconciliation and the costs of aggression in wild Barbary macaques (Macaca sylvanus): a test of the integrated hypothesis. Ethology. 2011 Oct;117(10):928–37.

41. Leone A, Mignini M, Mancini G, Palagi E. Aggression does not increase friendly contacts among bystanders in geladas (Theropithecus gelada). Primates. 2010 Oct;51(4):299–305.

